# Exploring the correspondence between gene expression and thalamic nuclei using the THALMANAC resource

**DOI:** 10.1101/2025.09.30.679413

**Authors:** Meghan A. Turner, Thomas Chartrand, Mathew T. Summers, Marcus Hooper, Cindy van Velthoven, Jack Waters, Saskia de Vries, Hongkui Zeng, Bosiljka Tasic, Karel Svoboda, Brian Long

**Author notes:** Equal contributors (Meghan A. Turner, Thomas Chartrand).

## Abstract

The thalamus connects the sensory organs and major subcortical brain regions with the neocortex. The thalamus has long been divided into multiple discrete nuclei, based on cytoarchitecture, histochemical stains, and mesoscale connectivity. However, thalamic nuclei do not completely describe thalamic organization. For example, some boundaries between thalamic nuclei are disputed, whereas other nuclei are known to contain subdomains with distinct connectivity and function. Moreover, the correspondence between cellular gene expression and other properties of thalamic projection neurons remains to be established. Spatial analysis of single cell gene expression provides a basis for reevaluating thalamic organization. We present the THALMANAC, (**THAL**amus**M**ERFISH **AN**alysis and **AC**cess) a Findable, Accessible, Interoperable, Reusable and Reproducible (FAIRR) resource for exploring and analyzing single-cell transcriptomic variation in the thalamus. The THALMANAC provides streamlined access to thalamic gene expression data registered to the common coordinate framework and tools for quantitative analysis and visualization of these data, all encapsulated in a reproducible, cloud computing platform. Using this resource, we find that gene expression generally supports the parcellation of thalamus into distinct nuclei. Some nuclei, such as the anteromedial nucleus, are additionally composed of discrete subdomains, while other nuclei share patterns of gene expression or are arrayed on a spatial gradient of gene expression. The THALMANAC establishes spatial transcriptomic data as a foundation for delineating thalamic organization.

**Graphical Abstract:** 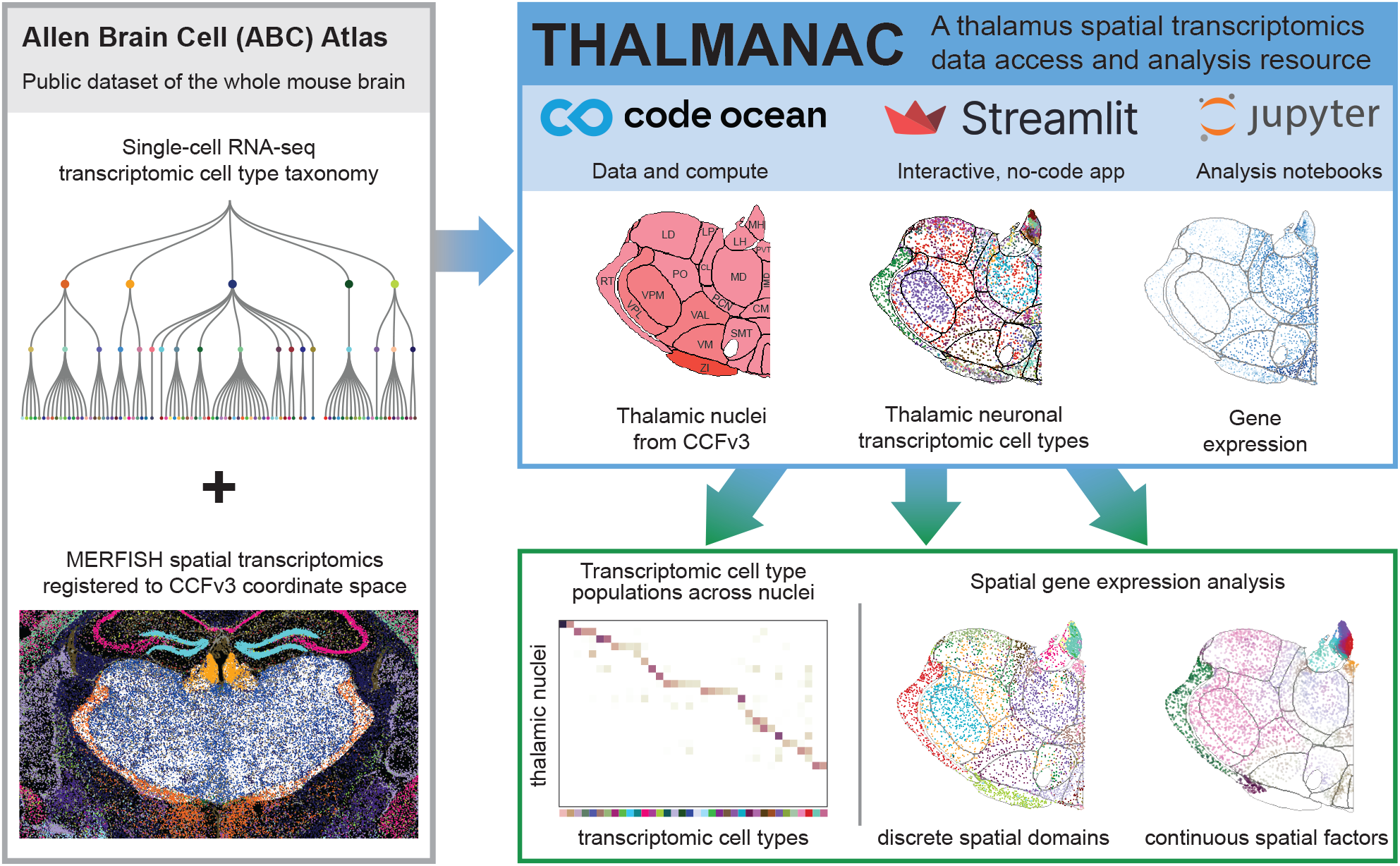

## Introduction

The thalamus is the central communication hub of the forebrain, relaying information from the sensory periphery, subcortical areas and the hippocampus to the cortex, in addition to mediating cortico-cortical communication (Jones 2007). The diverse inputs to and outputs from the thalamus are spatially organized (Harris et al. 2019; Hunnicutt et al. 2014). However, apart from the well-studied sensory thalamic nuclei, our understanding of thalamic organization in the context of brain-wide neural circuits is incomplete.

Anatomists have parcellated the thalamus into approximately 40 discrete nuclei, defined by combinations of cytoarchitecture (sizes, shapes, and density of cells; myelination patterns), histochemical stains, and mesoscale connectivity (Jones 2007). The parcellation of the mouse thalamus has been formalized in two widely used reference atlases: the 3D Brain Common Coordinate Framework (CCFv3) (Wang et al. 2020) and the 2D Mouse Brain in Stereotaxic Coordinates (also known as the Franklin-Paxinos Atlas, FP) (Paxinos & Franklin 2001; Paxinos & Franklin 2012). However, the atlases disagree on the shape and boundaries of many thalamic nuclei and the existence of many sub-nuclei. For example, the ventral posteromedial nucleus (VPM) is split into dorsal and ventral parts in FP (Chon et al. 2019), but not in the CCFv3. Moreover, recent anatomical and functional studies have shown that existing parcellations are not sufficiently detailed to capture known thalamic circuit function and organization (K. Guo et al. 2018; Z. V. Guo et al. 2017; Lee et al. 2020; Phillips et al. 2019). For example, distinct parts of VM form separate circuits with primary and secondary motor cortex (K. Guo et al. 2018). This fine-scale structure could correspond to discrete, molecularly defined subdivisions of VM that have not yet been annotated, or to topographic organization of thalamocortical and corticothalamic connectivity.

A major question is how gene expression relates to existing parcellations and thalamic organization in general. Single cell RNA sequencing (scRNA-seq) provides detailed measurements of transcription in large populations of individual neurons (Tasic et al. 2016), which can be used to define transcriptomic cell types (‘t-types’). The combination of scRNA-seq with spatial transcriptomics has been used to localize t-types across the whole mouse brain (Langlieb et al. 2023; Yao et al. 2023; Zhang et al. 2023). With t-types mapped into the CCFv3 coordinate space (Wang et al. 2020), we can evaluate how their spatial distributions correspond to anatomical parcellations (Yao et al. 2023).

Here, building on the Allen Brain Cell (ABC) Atlas scRNA-seq and spatial transcriptomics data, we developed a resource, the **THAL**amus **M**ERFISH **AN**alysis and **AC**cess (THALMANAC), which provides access to spatial gene expression data in the mouse thalamus; targeted analysis and data visualization of gene expression patterns; a platform for reproducible computations; and a no-code visualization app for data exploration. We demonstrate the utility of the THALMANAC by addressing how thalamic structural organization relates to gene expression. Aligning spatial transcriptomic data and nuclei boundaries reveals a wide range of relationships between t-types and thalamic nuclei: some t-types correspond to a single thalamic nucleus; others subdivide nuclei into multiple parts; many t-types span multiple thalamic nuclei. Analysis of expression by itself supports these same conclusions: some nuclei boundaries are associated with sharp changes in gene expression; other nuclei contain gene expression gradients; and, in many cases, gene expression patterns span multiple nuclei.

Understanding how information is routed through the thalamus requires integrating insights gained from gene expression with other modalities, including connectivity and neurophysiology. The THALMANAC provides a platform for incorporating additional spatial gene expression data to advance our understanding of thalamic organization.

## Results

### THALMANAC: A FAIRR resource for the thalamus

The THALMANAC resource provides access to thalamic spatial gene expression data, visualization of the relationships between gene expression, t-types, and CCFv3 anatomical parcellations, and quantitative analysis of targeted quantitative analyses, all encapsulated in a cloud platform that enables reproducible computations and customized exploration of the data.

The THALMANAC is built on a thalamus-specific subset of spatial transcriptomics data from multiplexed error-robust fluorescence *in situ* hybridization (MERFISH) from the ABC Atlas. The ABC Atlas dataset measures 500 transcripts, chosen to distinguish t-types across the whole mouse brain, in spatially resolved, single cells. MERFISH cells were registered to the CCFv3 coordinate space and mapped to the scRNA-seq data by assigning each MERFISH cell to the label of the most similar t-type cluster. We identified neurons that belong to the thalamus (including the zona incerta and habenula) according to their t-type label (**Figure S1A**) and CCFv3 parcellation label (**Figure S1B**). The thalamic dataset analyzed here includes 80,170 MERFISH cells across 14 coronal sections collected from a single male mouse (**Methods**).

The THALMANAC includes three avenues for engaging with the t-type and gene expression data and accompanying analyses (**Figure 1**). First, a Streamlit web app (RRID:SCR_024354) provides no-code exploration, including interactive visualizations and on-the-fly differential gene expression analysis for thalamic nuclei (Interactive Streamlit App: thalmanac.allenneuraldynamics.org). Second, Jupyter notebooks (RRID:SCR_018315) provide step-by-step demonstrations of the Python code needed to reproduce all results presented here (github.com/AllenNeuralDynamics/thalmanac). Both tools allow, users to customize the thalamic cells that are selected from the full ABC Atlas. Users can also choose to visualize alternative anatomical registrations: a section-wise affine alignment to CCFv3 (**Methods**) and an alignment to the DevCFF atlas (Kronman et al. 2023). Third, an AnnData object (Virshup et al. 2024) containing the thalamic spatial transcriptomics subset of the ABC Atlas and our *de novo* analysis results is accessible via the python API or local download and is suitable for further exploration in widely used visualization tools such as Cirrocumulus (RRID:SCR_021646) and ManiVault Studio (Vieth et al. 2024).

**Figure 1.**
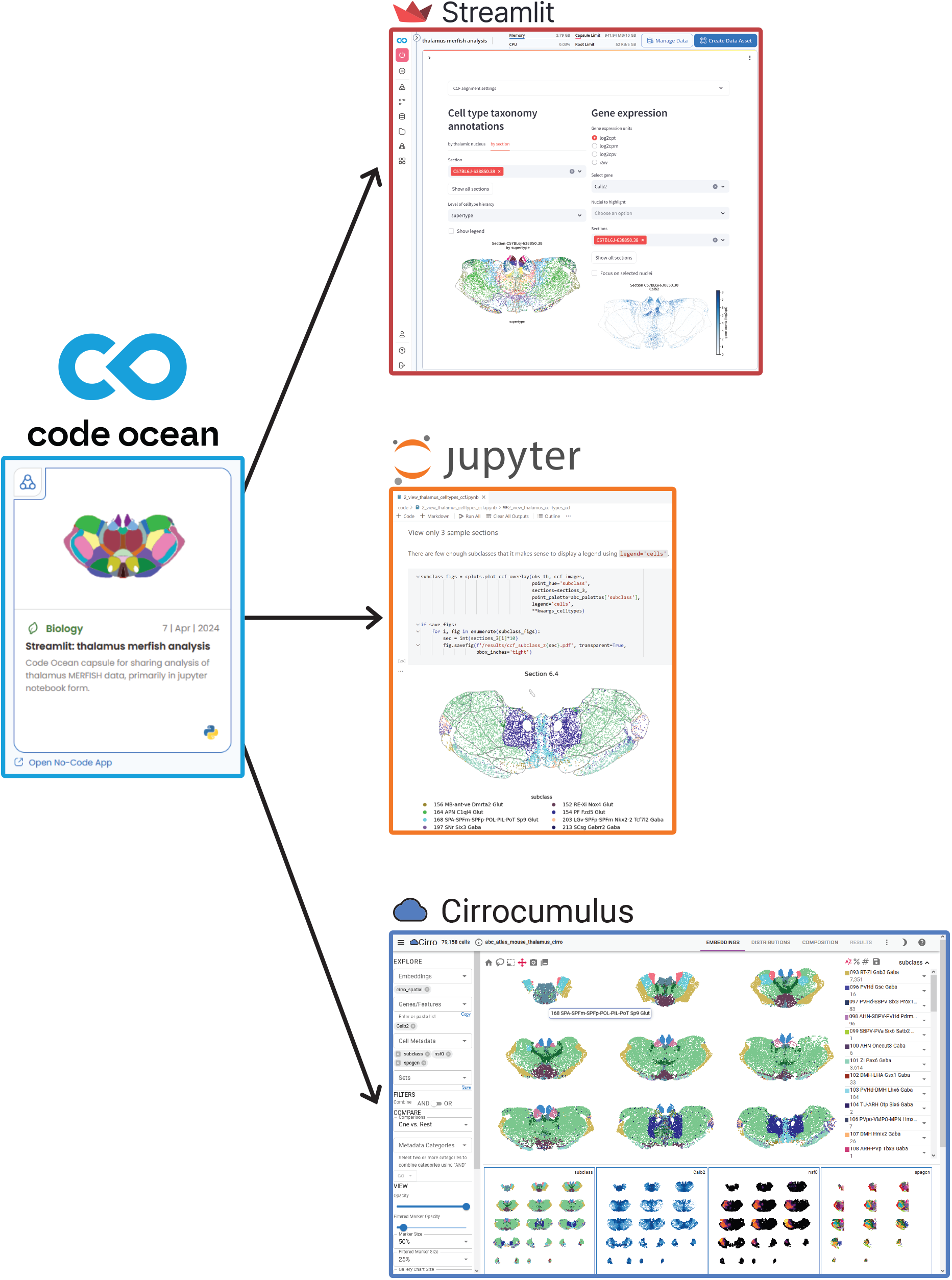
Interactive and customizable visualization tools for exploration of the thalamus MERFISH dataset hosted on the Code Ocean cloud platform. Diagram illustrating the three workflows that can be launched from the Code Ocean Collection accompanying this paper. The Code Ocean platform provides built-in support for code versioning (via Git), environment replicability (via Docker), data and analysis provenance (via tracked data assets and linked pipeline runs), cloud compute resources (via AWS EC2 instances). Users can launch an interactive, no-code Streamlit application to explore spatial distributions of transcriptomic cell types and gene expression from their browser, as well as visualize locations of annotated cell types for each thalamic nucleus and highlight a specific nucleus or group of related thalamic nuclei. Tutorial Jupyter notebooks and Python code replicate all analyses and visualizations in a cloud workstation. These visualizations can be customized and extended. Users can download an AnnData-formatted .h5ad file (RRID:SCR_018209), which is compatible with widely used open source software, such as Cirrocumulus (RRID:SCR_021646).

All components of the THALMANAC are hosted on the Code Ocean platform (Code Ocean Collection). Code Ocean provides a cloud-based solution for the reproducibility challenge in the analysis of large biological datasets (Meijer et al. 2025), such as those generated by spatial transcriptomics platforms. A self-contained Code Ocean ‘capsule’ (2025) has built-in management for each of the FAIRR (Findable, Accessible, Interoperable, Reusable, Reproducible) principles (Koster & Liu 2024; Wilkinson et al. 2016), as well as all features required to reproduce the analyses described here, including: computing environment via Docker container (RRID:SCR_016445); data provenance via a tracked data asset that links directly to the cloud-hosted ABC Atlas data (RRID:SCR_024440), code versioning via Git (RRID:SCR_003932), and preconfigured cloud compute resources.

In the following sections, we describe how we used the THALMANAC to explore the relationships between gene expression, t-types and thalamic nuclei.

### Correspondence between thalamic nuclei and transcriptomic types

In the ABC Atlas, transcriptomic variation is used to assign cells to a hierarchy of discrete t-types (from broad to narrow categories: class, subclass, supertype, cluster). Thalamic neurons belong to 5 *classes*, 58 *subclasses*, 163 *supertypes*, and 563 *clusters* (**Table S1**; **Figure S2**) (taxonomy version **CCN20230722**) (Yao et al. 2023), but these numbers will change as the taxonomy evolves. Whereas the cluster level is based on unsupervised clustering, higher levels of the taxonomy are additionally informed by developmental origin via shared transcription factor expression, anatomical location, and classical (pre-scRNAseq) neuron types (DeFelipe et al. 2013) defined by key molecular markers (e.g. neuropeptides, calcium-binding proteins).

Transcriptomic *classes* typically correspond to major brain structures. Glutamatergic neurons in thalamus proper form a single class (class 18 TH Glut), whereas habenula neurons belong to a distinct class (class 17 MH-LH Glut) (**Figure S2B**). The other three classes are not unique to the thalamus: class 12 HY GABA is found in the reticular thalamus, zona incerta and ventral lateral geniculate nucleus (CCFv3 annotations; RT, ZI, and LGv, respectively, see **Table S1**) but also in the hypothalamus; class 19 MB Glut is found in thalamic nucleus SPA but also in the midbrain; class 20 MB GABA is found in thalamic nuclei LGv and SPF and the midbrain. Transcriptomic *subclasses* span multiple thalamic nuclei(**Figure S2C**), whereas transcriptomic *supertypes* often define individual nuclei (e.g. AM), and several nuclei contain multiple supertypes (e.g. ZI) (**Figure S2D**). Most nuclei contain multiple transcriptomic *clusters* (**Figure 2A, Figure S2E**).

**Figure 2.**
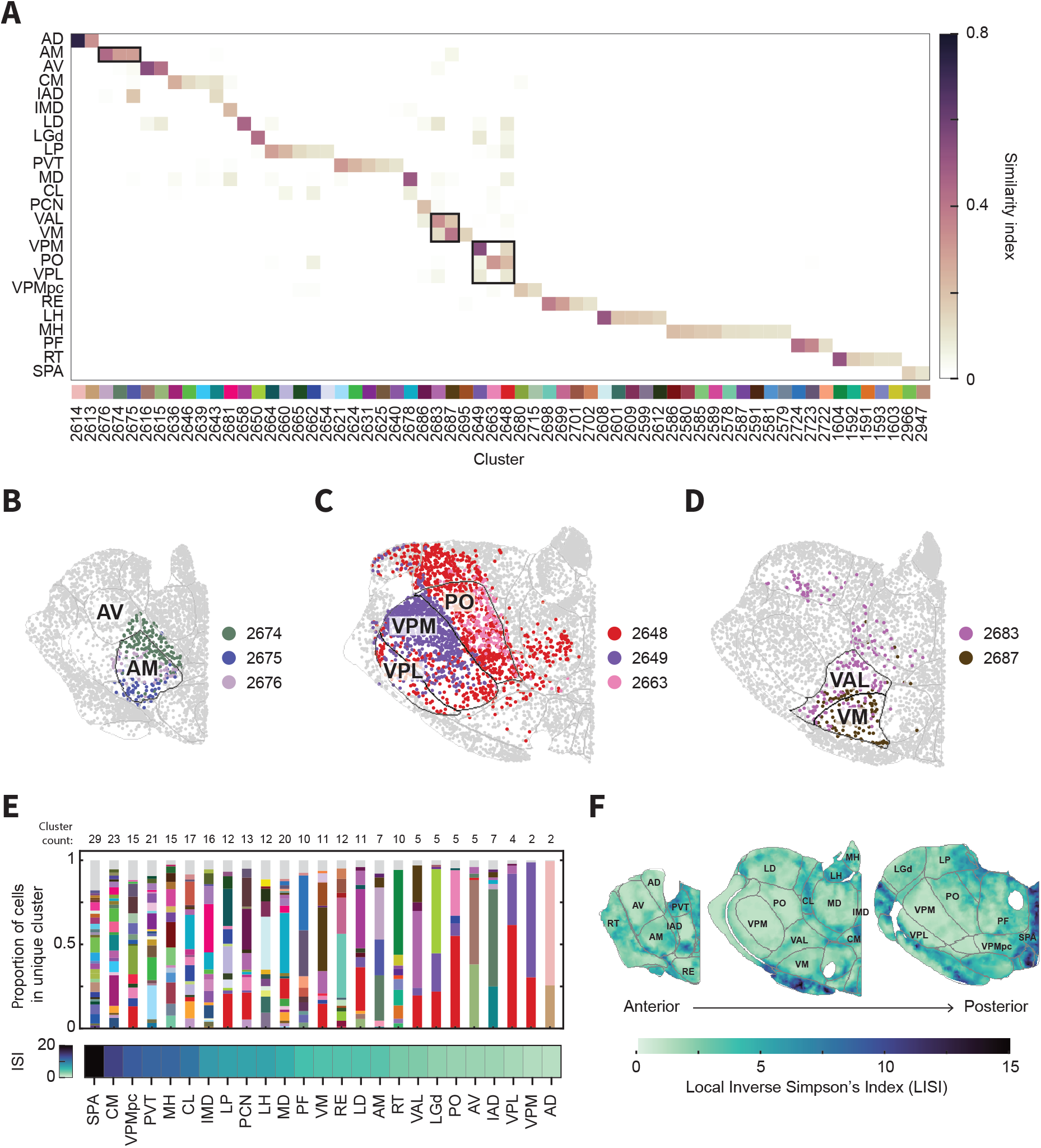
Transcriptomic diversity across the thalamus. **(A)** Similarity of the boundaries of thalamic nuclei to the spatial distribution of transcriptomic clusters (**Methods**). Thalamic nuclei names and abbreviations are listed in **Table S2** (asterisks in the table indicate the 25 nuclei that are included in this figure). **(B)** Clusters that primarily overlap the anteromedial (AM) nucleus (section C57BL6J-638850.43). **(C)** Clusters that overlap with the somatosensory nuclei (VPM, VPL, and PO) (section C57BL6J-638850.37). **(D)** Clusters that primarily overlap the ventromedial (VM) and ventro-anterior-lateral (VAL) nuclei (section C57BL6J-638850.39). **(E)** The proportion of cells in specific clusters for each thalamic nucleus. Thalamic nuclei are ordered by their Inverse Simpson’s Index (ISI, bottom). Only clusters that represent > 1% of cells in a nucleus are given a unique color in the barplot and included in the cluster count numbers. Clusters with <1% of cells are grouped in “other” (grey). **(F)** Spatial map of the local cluster diversity across the thalamus, as quantified by the inverse Simpson’s index (ISI) of the clusters found in each cell’s 15 nearest neighbors. The 15 nearest neighbors for the local diversity calculation are restricted to cells found in the same MERFISH section (sections, left to right: C57BL6J-638850.44, C57BL6J-638850.40, C57BL6J-638850.36).

We explored correspondence between clusters and thalamic nuclei. We quantified the match between cluster and nucleus as a similarity index based on cell-wise overlap (**Figure 2A**; **Figure S3**; **Methods**). We annotated the best nucleus-cluster matches, starting with pairs of similarity greater than 0.1. We then refined initial annotations by a semi-automated process to remove spurious matches and ensure at least a single match for each cluster and nucleus (**Methods, Table S3**).

The correspondence between the boundaries of thalamic nuclei and the spatial distributions of clusters varies. In some nuclei, such as AM, the spatial distribution of the matched clusters aligns to the nucleus boundary (**Figure 2B**; **Figure 3**). Other nuclei, such as VPM, have a single strongly matched cluster, with another cluster that spills beyond the VPM boundary (**Figure 2C**; **Figure 4**). In other cases, the matched clusters are shared between a few nuclei (**Figure 2D**). VM primarily contains cluster 2687 and VAL primarily contains cluster 2683, but both clusters cross the nucleus boundaries, showing a gradual transition of populations from VAL to VM.

**Figure 3.**
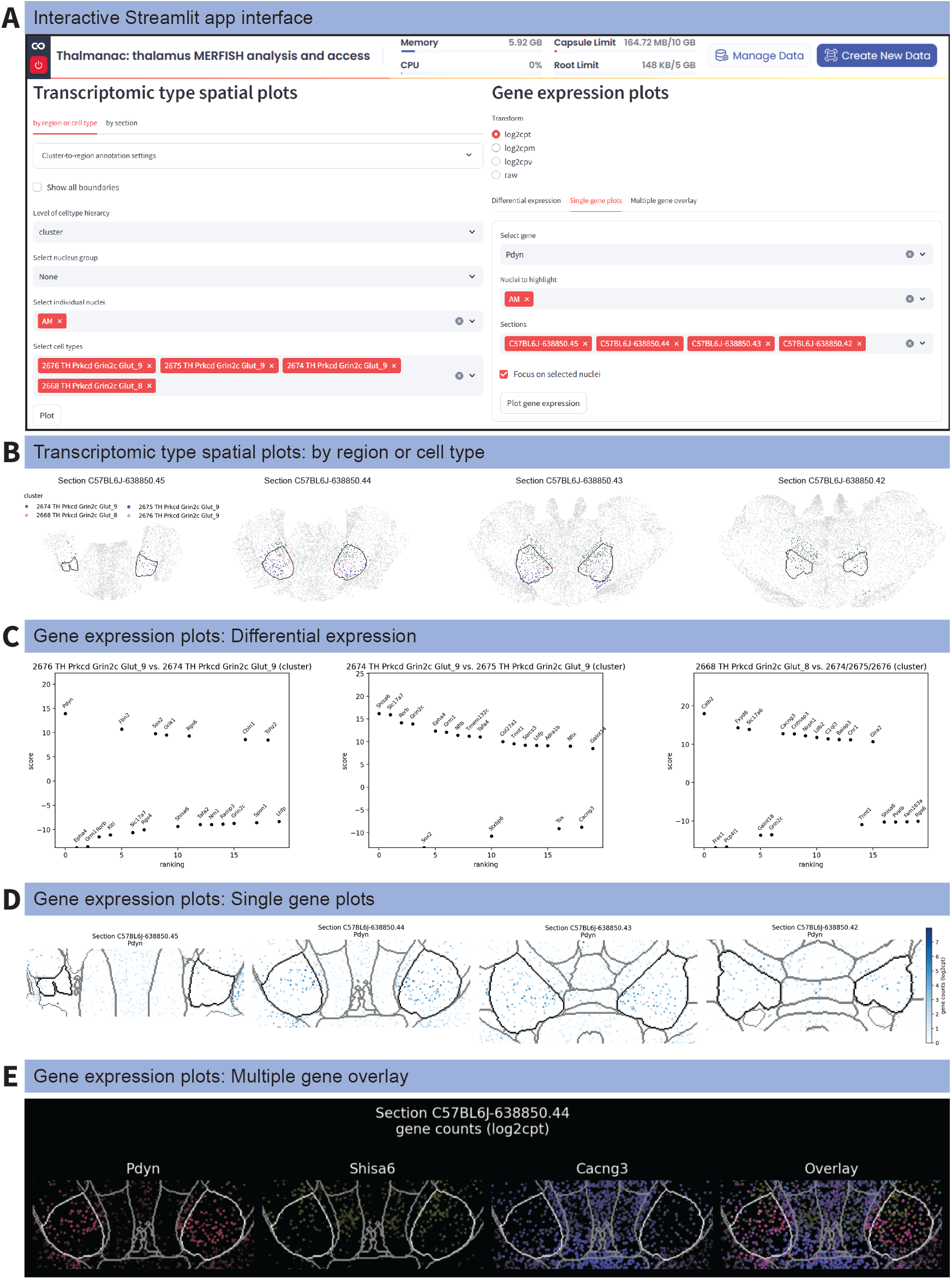
Spatially variable genes distinguish adjacent transcriptomic clusters in the AM nucleus. **(A)** No-code Streamlit application interface with plotting options selected for the AM (anteromedial) thalamic nucleus. Lettered icons indicate which application panels have their outputs shown in the other figure panels. **(B-E)** Output figures generated from the Streamlit application using the settings indicated in (A). **(B)** *Transcriptomic type spatial plots – by region or cell type*: Spatial locations of the four transcriptomic clusters found in AM: cluster 2674, 2675, 2676, and 2668 (sections C57BL6J-638850.45, C57BL6J-638850.44, C57BL6J-638850.43, and C57BL6J-638850.42). **(C)** *Gene expression plots – DiWerential expression:* Top twenty differentially expressed gene rankings for pairs of AM clusters (left to right: cluster 2676 vs 2674, 2674 vs 7675, and 2675 vs 2676). **(D)** *Gene expression plots – Single gene plots*: Spatial gene expression pattern for the top differentially expressed gene in cluster 2676, *Pdyn*. Plots were focused on the AM nucleus by checking the Focus on selected nuclei box as shown in (A). **(E)** *Gene expression plots – Multiple gene overlay*: Spatial gene expression pattern for the top differentially expressed gene from each cluster pair in (C): *Pdyn, Shisa6*, and *Calb2*. Genes shown individually and as a multi-channel overlay (section C57BL6J-638850.44). Plots were focused on the AM nucleus by checking the *Focus on selected nuclei* box as shown in (A).

**Figure 4.**
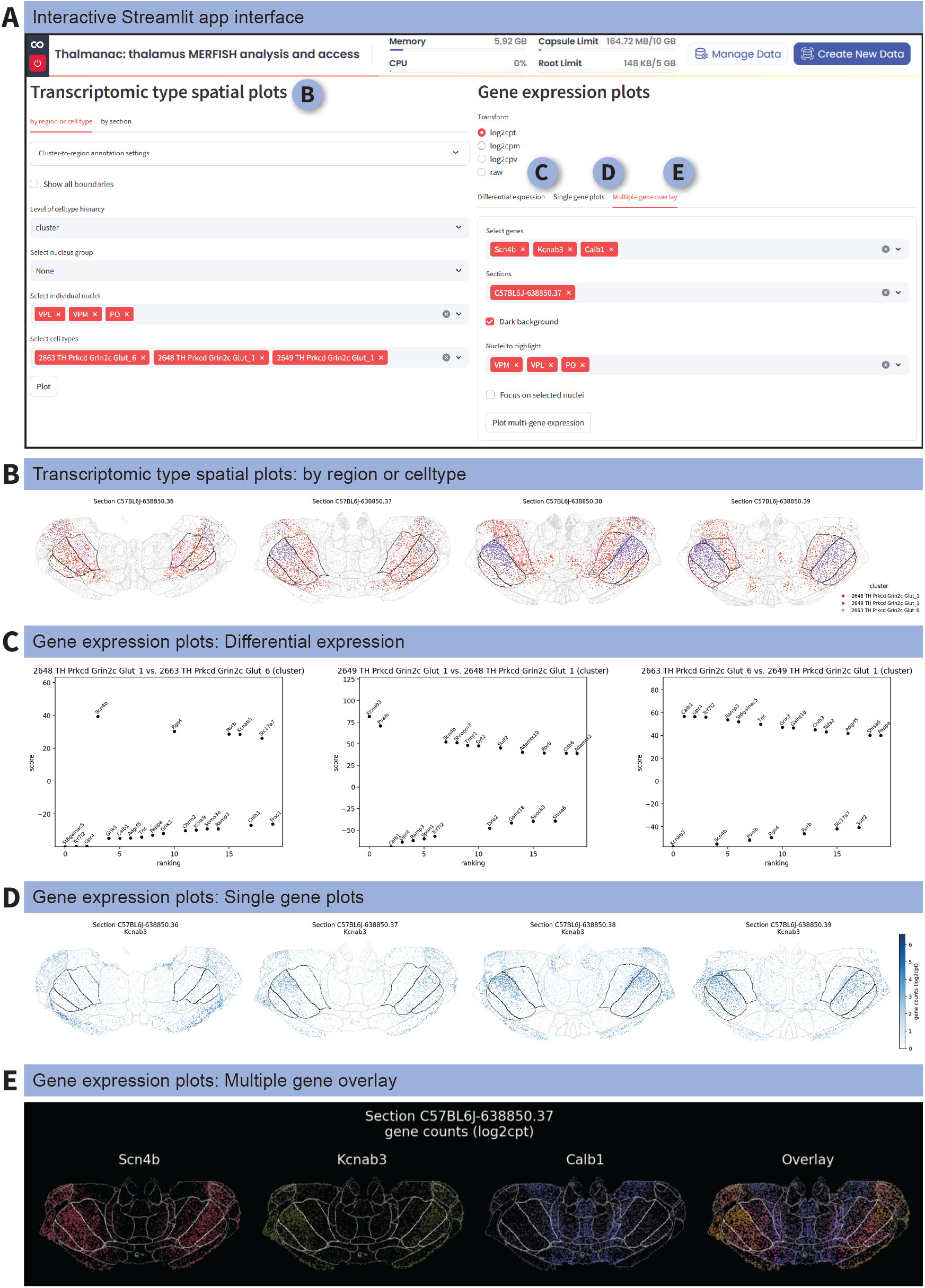
Transcriptomic clusters and spatially variable genes are shared across the somatosensory nuclei. **(A)** No-code Streamlit application interface with plotting options selected for the somatosensory nuclei of the thalamus: VPM, VPL, and PO. Lettered icons indicate which application panels have their outputs shown in the remaining figure panels. **(B-E)** Output figures generated from the Streamlit application using the settings indicated in (A). **(B)** *Transcriptomic type spatial plots – by region or cell type*: Spatial locations of three transcriptomic clusters found in the somatosensory nuclei and neighboring nuclei: clusters 2648, 2649, and 2663 (sections C57BL6J-638850.36, C57BL6J-638850.37, C57BL6J-638850.38, and C57BL6J-638850.39). The CCFv3 structures of VPM, VPL, and PO are outlined in black. **(C)** *Gene expression plots – DiWerential expression:* Top twenty differentially expressed gene rankings (top 20 genes shown) for pairs of somatosensory transcriptomic clusters (left to right: cluster 2648 vs 2663, 2649 vs 2648, and 2663 vs 2649). **(D)** *Gene expression plots – Single gene plots*: Spatial gene expression for pattern for the top differentially expressed gene in cluster 2648, *Kcnab3*. **(E)** *Gene expression plots – Multiple gene overlay*: Spatial gene expression pattern for the top differentially expressed gene from each cluster pair in (C): *Scn4b, Kcnab3*, and *Calb1*. Genes shown individually and as a multi-channel overlay (section C57BL6J-638850.37). Plots were focused on the somatosensory nuclei by checking the Focus on selected nuclei box as shown in (A).

We quantified the range of cluster-level diversity found in different thalamic nuclei in three ways (**Figure 2E-F**). First, we calculated the cluster count, which is the number of clusters present in each nucleus (**Figure 2E**). SPA contains the largest number of clusters, followed by CM, PVT, and MD; AD and VPM contain the fewest clusters.

Second, we calculated the Inverse Simpson’s Index (ISI), in which the cluster counts are weighted by cluster size to account for the relative proportions of clusters in each nucleus (**Figure 2E**). High ISI values highlight nuclei whose cells are distributed amongst many clusters, such as SPA, CM, VPMpc, and PVT. In contrast, low values identify nuclei dominated by few clusters, such as AD, VPM, VPL, and IAD. Some nuclei, such as MD, rank high in cluster count but rank lower in ISI, indicating that they are dominated by a subset of their clusters.

Third, to identify sub-structure within a nucleus, we calculated the Local Inverse Simpson’s Index (LISI), which is the ISI of each cell and its 15 nearest neighbors (**Figure 2F**). We found uneven cluster density, especially within nuclei that contains many clusters, such as SPA and CM. As expected, cluster diversity is higher at the boundaries between nuclei, especially between neighboring nuclei that contain distinct transcriptomic types, such as AV and AM.

Interpreting these metrics together allows us to integrate different facets of transcriptomic diversity into a single picture for each nucleus. For example, the cells in SPA are distributed relatively evenly (high ISI) amongst its many clusters (high cluster count of 29). However, all but the two clusters with a high similarity index are not unique to SPA. And these clusters do not form distinct spatial domains within SPA but instead are mixed together (high LISI).

### Spatially variable gene expression within and between nuclei

The spatial patterns of t-types can be dominated by large, discrete, gene expression differences in one or a few genes or by smaller, continuous, gene expression differences in a larger numbers of genes. THALMANAC facilitates exploration of the spatial gene expression patterns that drive the relationship between t-types and thalamic nuclei. We provide examples of this exploration using the no-code Streamlit application (**Figure 3A, Figure 4A**). Based on cluster-nucleus annotations, we select four transcriptomic clusters corresponding to the anterior medial thalamic nucleus (AM) (**Figure 3B**). These clusters are found only in and around the AM in a complex spatial pattern. We use the differential gene expression analysis feature in the app to identify the gene expression patterns that underlie these clusters (**Figure 3C**). The central AM cluster, 2676, can be distinguished by its high expression of *Pdyn*, which is minimally expressed in the two outer clusters, 2674 and 2675 (**Figure 3C,D**). *Shisa6* is preferentially expressed in the most dorsomedial cluster, 2674, and minimally expressed in the most ventrolateral cluster, 2675 (**Figure 3C,E**). Conversely, *Cacng3* is preferentially expressed in the most ventrolateral cluster, 2675, and minimally expressed in the most dorsomedial clusters, 2674 and 2676(**Figure 3C,E**). This spatial variation of gene expression within AM is consistent with recent spatial transcriptomic work that focused specifically on the AM and neighboring thalamic nuclei (Kapustina et al. 2024).

We also use the THALMANAC to explore a group of nuclei, the somatosensory nuclei (VPM, VPL, and PO) (**Figure 4A**). In contrast to AM, the somatosensory nuclei share three t-types with each other and with other neighboring nuclei (**Figure 4B**). Cluster 2649 is shared between VPL, PO, and the antero-ventral portion of VPM, as well as neighboring nuclei such as LD and VM. Cluster 2648 is primarily localized to the postero-lateral portion of VPM and cluster 2663 is mostly localized to PO. Differential gene expression analysis (**Figure 4C**) highlights some of the spatially variable gene expression gradients that underly these shared t-types (**Figure 4D,E**). *Kcnab3* is more highly expressed in cluster 2649 but is also expressed at lower levels in cluster 2648 (**Figure 4D**). This suggests that these two clusters lie on a gene expression gradient, rather than a sharp boundary of gene expression. The limited transcriptomic diversity identified by this dataset does not fully capture known functional and anatomical topography (e.g. barreloids in the VPM corresponding to individual whiskers (Van Der Loos 1976)). The somatosensory nuclei highlight the need for multi-modal datasets—combining transcriptomic, functional, and connectivity data—to fully understand the principles of thalamic organization.

### Unsupervised spatial patterns of thalamic gene expression

The preceding approaches analyzed correspondence between gene expression and thalamic nuclei based on t-types or expression of single genes. Other patterns relevant to the spatial organization of thalamus may involve combinations of many genes. Most transcriptomic types in the taxonomy are defined by multi-gene patterns, but without spatial context. Searching for patterns in spatial gene expression *de novo* (without reference to taxonomy or nuclear divisions) has the potential to discover new organization across the thalamus and to provide confirmation for spatial patterns highlighted by t-types. We applied two unsupervised spatial analysis methods to the MERFISH gene expression data, representative of distinct approaches to pattern discovery. Domain detection methods, such as the SpaGCN package (Hu et al. 2021), assign each cell to a single pattern to produce spatially contiguous regions or domains. In contrast, factor methods, such as non-negative spatial factorization (NSF) (Townes & Engelhardt 2023), identify graded patterns that resemble gene expression programs and allow multiple patterns to contribute to each cell’s identity.

We applied both methods to identify sets of 30 distinct patterns, approximately corresponding to the number, or level of resolution, of thalamic nuclei. Using SpaGCN, we found that the discrete domains identified had strong overlap with classically defined thalamic nuclei (**Figure 5A-C**). For example, domains 25, 17, and 16 align with the outer boundaries of AD, AV, and AM, respectively, and domain 0 aligns with the boundaries of RT (**Figure 5B**). In comparing this *de novo* domain detection to the scRNA-seq transcriptomic types, we find that the domains most often resemble organization at the subclass or supertype levels of the taxonomic hierarchy, rarely picking up on the finest cluster level distinctions identified by mapping spatial cells to the RNA-seq taxonomy (**Figure S4**).

**Figure 5.**
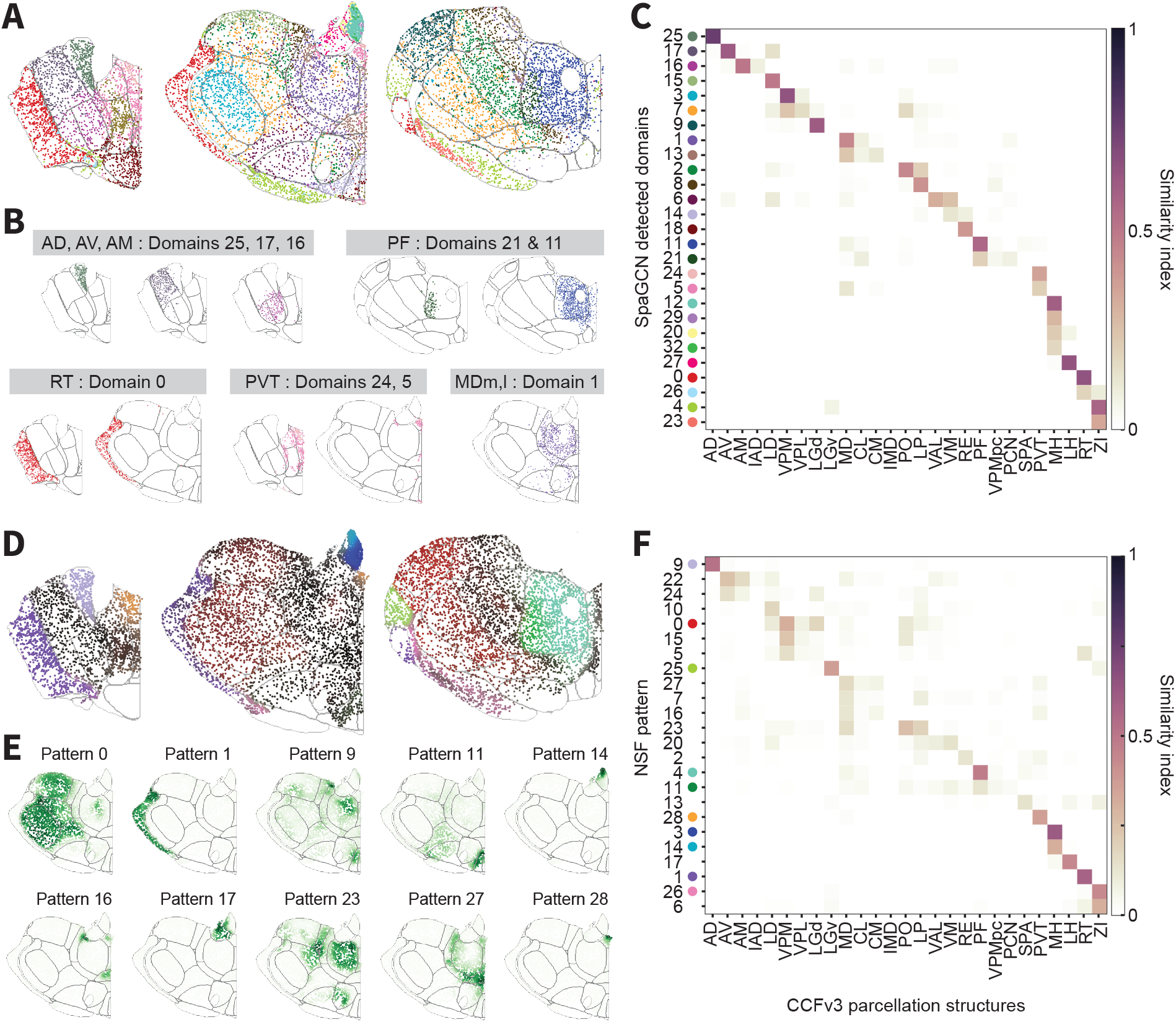
Unsupervised spatial clustering reveals both discrete & continuous gene expression patterns in the thalamus. **(A)** Domains detected by the SpaGCN algorithm (sections, left to right: C57BL6J-638850.44, C57BL6J-638850.40, C57BL6J-638850.36). Colored markers and domain identifiers along the y-axis of (C) serve as a color legend. **(B)** Spatial locations of nine SpaGCN domains that closely correspond to seven thalamic nuclei. **(C)** Heatmap quantifying the similarity in spatial distribution of SpaGCN domains to the anatomically defined boundaries of thalamic nuclei. Colored markers next to the domain identifiers (y-axis) serve as a color legend for (A). **(D)** Overlay of 10 spatial patterns (factors) generated by the non-negative spatial factorization (NSF) algorithm in three example sections across the anterior-posterior axis. Colored markers and pattern identifiers along the y-axis of (F) serve as a color legend. **(E)** Individual plots of ten spatial patterns from (D). **(F)** Heatmap quantifying the similarity in spatial distribution of SpaGCN domains to the anatomically defined boundaries of thalamic nuclei. Colored markers next to the pattern identifiers (y-axis) serve as a color legend for (D). For heatmaps in (C) & (F), thalamic nuclei names and abbreviations included in heatmaps are listed in **Table S2.** Similarity was measured using the Dice coefficient (see **Methods**).

The NSF method identifies sparse, partially overlapping programs of gene expression across and within thalamic nuclei boundaries. Some individual NSF patterns strongly matched single thalamic regions like RT or MH, indicating distinct and coherent gene expression programs. Many other patterns span neighboring nucleus boundaries or highlight connections between distant nuclei, such as pattern 0 (spanning VPM, PO, LD, and MDc) or pattern 23 (spanning LP, PO, MD, and SMT). Single genes contributing to each pattern can also be identified, and in some cases can nearly reproduce the pattern alone, providing markers for this graded view of thalamic transcriptomic variation (**Figure S5**).

## Discussion

A comprehensive understanding of how the thalamus routes information to the cortex will require multiple types of data and a community-wide effort. Our work contributes to this program by curating and analyzing the correspondence between state-of-the-art spatially resolved, single-cell transcriptomic data, and widely used parcellations of thalamus into discrete nuclei. In addition, we make our data and analyses available in an accessible, documented, and reproducible bioinformatics resource in the cloud. Using the AM nucleus and the somatosensory nuclei as examples, we illustrate how this interactive resource can be used to identify novel patterns of gene expression and how these patterns correspond to prior parcellations of the thalamus.

The THALMANAC resource streamlines researchers’ access to thalamic spatial transcriptomic data. We anticipate that scientists will extend our analysis based on the ABC atlas data, or apply similar analyses to additional spatial transcriptomics datasets registered to the CCF coordinate system(Langlieb et al. 2023; Zhang et al. 2023). The tool should also be used as a reference to validate recent studies focused on specific nuclei, such as the spatially segregated projections from PF to distinct dorsal striatum regions (Mandelbaum et al. 2019) and spatially segregated inputs to PO (Casas-Torremocha et al. 2022), the molecular organization of the PVT (Gao et al. 2023), and the core/shell organization of neurons in RT (Li et al. 2020). Our interactive Streamlit app can be used to identify marker genes for post-hoc localization of neurophysiological and neuroanatomical experiments, or to select gene panels for multiplexed mFISH experiments. The THALMANAC is already being used in the effort to create and characterize enhancer-based genetic tools (Ben-Simon et al. 2025) to target thalamic cell types and regions.

The THALMANAC shows that at the scale of thalamic nuclei, the molecular organization of the thalamus resembles existing parcellations. However, both discrete cell type composition and gene expression gradients vary at finer spatial scales within and across nuclei. Given the evidence for circuit and functional variation at the same length scales, additional data need to be integrated with molecular characterization to understand thalamic organization. We see two promising paths forward for data integration. First, existing data registered to the CCFv3 3D coordinate space, such as the databases of single-neuron morphological reconstruction (Winnubst et al. 2019) or neurophysiology data (Chen et al. 2024), could be linked to cell types and gene expression described in the THALMANAC. Second, targeted multimodal datasets, such as single-neuron morphological reconstructions combined with *post-hoc* mFISH for transcriptomic cell typing would link cell types and gene expression to projection targets at the single cell level.

Beyond data integration, we also recognize the need for thalamus-focused, community annotation, similar to previous efforts to annotate the fly connectome (Schlegel et al. 2024). We envision a refined parcellation of the thalamus where scientists with specific domain knowledge would combine molecular information from the THALMANAC with neuroanatomical and functional data to reflect progress in understanding of thalamic organization.

## Methods

### ABC Atlas mouse whole-brain MERFISH dataset

The four million cell MERFISH dataset from the public Allen Brain Cell (ABC) Atlas mouse whole-brain transcriptomic cell type atlas (Yao et al. 2023) was downloaded as instructed by the ABC Atlas access documents.

### Thalamus dataset

To subset the published ABC Atlas MERFISH dataset (3,739,961 cells from 59 coronal sections) to just those cells that belong to the thalamus (80,170 neurons from 14 coronal sections), we filtered on both anatomical region and transcriptomic types. To filter by anatomical region, we used the ABC Atlas’ rasterized anatomical annotation image volume, which aligns the Allen CCFv3 3D coordinate space, resampled at 10um voxel resolution, into the MERFISH data space. In this image volume, each voxel is assigned a *parcellation_index* that uniquely identifies its assigned Allen CCFv3 annotation (RRID:SCR_020999), as described in the ABC Atlas tutorial notebooks. We selected all voxels labelled with a TH-or a ZI-associated parcellation_index (**Figure S1A(ii)**; **Table S2**) to generate a binary mask of the thalamus (**Figure S1A(iii)**). To capture any TH or ZI cells that fall outside the averaged CCFv3 structure boundaries, we first expanded the binary mask, using a binary closing operation (`scipy.ndimage.binary_closing`) followed by a binary fill holes operation (`scipy.ndimage.binary_fill_holes`), to include any white matter tracts inside the boundaries of the thalamus and then dilated the binary mask by 20um to include cell just outside the exterior TH boundary (**Figure S1A(iv)**; **Methods**). Furthermore, to remove a detached mask region from a posterior section, we removed any binary mask regions whose area ratio, as compared to the largest binary mask region in that same section, was <0.1. We filtered out any cells that fell outside this final binary mask volume (**Figure S1B(ii)**). This reduced the dataset to 162,117 cells and 16 coronal sections.

Finally, upon visual inspection, we filtered out all cells from the anterior-most section and the posterior-most section that contained TH and ZI cells (C57BL6J-638850.46 and C57BL6J-638850.29, respectively) due to poor alignment between the binary mask derived from the CCFv3 annotations and cells mapping to thalamic transcriptomic types. This further reduced the dataset to 160,663 cells and 14 coronal sections.

To further curate the dataset according to mapped scRNA-seq transcriptomic type, we first filtered out cells that belonged to the four non-neuronal classes: “30 Astro-Epen”, “31 OPC-Oligo”, “33 Vascular”, and “34 Immune” (**Figure S1B(iii)**). We then manually determined that nearly all neurons in the TH and ZI (94%) belonged to three thalamus-associated neuronal classes (“12 HY GABA”, “17 MH-LH Glut”, and “18 TH Glut”), while a small percentage of cells in the TH—including nearly all neurons in the SPA and a sparse population of interneurons in the LGd—mapped to two midbrain-associated neuronal classes (“19 MB Glut”, 3.5%, and “20 MB GABA”, 2.5%). We filtered out any cells that did not belong to these five “thalamic neuronal” classes (**Figure S1B(iv)**). This resulted in the final thalamus dataset containing 80,170 neurons across 14 coronal sections.

Corresponding code for the steps described above can be found in the Jupyter notebook 7_sub-set_abc_atlas_to_thalamus.ipynb. We demonstrate how to use the thalamus_merfish_analysis package to: (1) load the standard thalamic neuronal dataset in the 1a_load_thalamus_data_standard Jupyter notebook; and (2) load custom or alternative thalamus subsets of the ABC Atlas in the 1b_load_thalamus_data_custom Jupyter notebook.

### Cell-wise similarity index

For each thalamic nucleus-cluster pair, the overlap between cells labeled as belonging to both the nucleus and the cluster was quantified using the Dice similarity index: the size of the set intersection (cells in both nucleus and cluster) divided by the mean of the nucleus and cluster sizes. A similarity index of 1 indicates complete overlap between cells in the nucleus and in the cluster, whereas a value of zero indicates no overlap between these two labels. The same similarity measure was used to quantify the overlap between cells labeled with the SpaGCN domain detection and cells belonging to each thalamic nucleus. For NSF patterns, this was generalized to the Quantitative Dice index (allowing each cell to match a distribution over multiple patterns rather than a single pattern), and calculated as 1-d, the Bray-Curtis dissimilarity (from sklearn’s ‘metrics.pairwise_distances’).

To account for misalignment and possible distortions of thalamic nuclei boundaries in the Allen CCFv3 parcellation (caused by the process of combining multiple individuals to generate an average template), we excluded any cell which fell within 50 µm of nucleus boundaries. This process minimized the impact of distortions at most nuclei boundaries, but we found examples where it was not sufficient to compensate for poor alignment.

### Thalamic nucleus-to-cluster annotations

An annotation of the best-matched clusters for each thalamic nucleus was created using the similarity index results (**Table S3**). Matches were annotated based on a cutoff of similarity>0.05 for the top match of each cluster or nucleus and similarity>0.1 for additional matches. Additionally, we manually refined the set of annotated clusters on a per-nucleus basis by visually inspecting the matches with nuclei boundaries from both alignments, removing clusters that appeared to be outliers or only present in misaligned regions, and adding relevant additional matches that were below the cutoff as needed for near-complete coverage of the nucleus.

### Diversity metrics

Cluster count was quantified as the number of unique clusters present in each nucleus. For **Figure 2B**, clusters were only included in the final count if they represented >1% of all cells in the nucleus; all clusters representing <1% of cells in a nucleus were grouped into an “other” category.

The Inverse Simpson’s Index (ISI), was defined as:

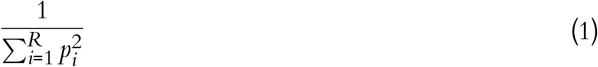

where R is the number of distinct clusters present and pi is the proportional abundance of each cluster.

The Local Inverse Simpson’s Index (LISI) applies the above ISI formula to each cell and its 15 nearest neighbors.

### Differential gene expression analysis

Differential expression analysis, as shown in **Figure 3** and **Figure 4**, was performed with scanpy’s `rank_genes_group` function. The top 20 ranked genes (positive or negative) were returned.

#### Domain detection with SpaGCN

Spatial domains were generated using the SpacGCN package (Hu et al. 2021) with the following parameters and modifications.

The parameter, *p*, which controls the percentage of total expression contributed by a cell’s neighborhood, was set to 0.5. The length decay parameter, *l*, was calculated from *p* for each section using SpaGCN’s provided `search_l` function and then averaged together to generate a global *l* parameter. All random seeds were set to 100. Clusters were initialized with Louvain clustering. We conducted a parameter sweep for resolutions of the Louvain clustering (1.4, 1.0, 0.7, 0.4). We manually selected a resolution of 1.4 for inclusion in this study, which detects 32 spatial domains, for its reasonable agreement with thalamic nuclei organization. Increasing the resolution beyond 1.4, and thus the number of detected spatial domains, only serves to increase the noise of the domains, resulting in salt-and-pepper patterns rather than finding additional, smaller contiguous domains as one might like.

To generate unified domains across all 14 thalamic sections, we calculated the cell-cell distance matrix for each section separately and then diagonal block concatenated those individual matrices together, filling off diagonal entries with infinite values. This results in a single, multi-section adjacency matrix where cells in different sections have an adjacency value of zero, as previously implemented in the domain detection package STAligner (Zhou et al. 2023). This additional code is available as part of our THALMANAC Code Ocean Collection.

### Non-negative spatial factorization

Gene patterns from non-negative spatial factorization were generated using the NSF package (Townes & Engelhardt 2023) with the following parameters and modifications. Key parameters included the number of patterns: 30; length scale: 200 µ*m*; number of inducing points (IPs): 2000; and batch size: 5000. Since the full model fitting process required more memory than available on a standard GPU, we added multi-GPU compatibility by subclassing the SpatialFactorization and ModelTrainer classes, adding appropriate TensorFlow decorators and minor updates to incompatible code. This additional code is available as part of our THALMANAC Code Ocean Collection.

## Data and code availability

All data and code described in this publication are publicly available. Code can be accessed either on it’s own via our GitHub repository (github.com/AllenNeuralDynamics/thalmanac) or alongside computational resources and the data via our THALMANAC Code Ocean Collection (codeocean.allenneuraldynamics.org/collections/a15ba7ef-bcce-4593-bacb-432bd8bf0596). Our interactive THALMANAC Streamlit app can be accessed at: thalmanac.allenneuraldynamics.org.

The dependency package for accessing and processing the public ABC Atlas MERSCOPE data can be accessed via an accompanying GitHub repository: github.com/AllenNeuralDynamics/abc-merfish-analysis.

## Acknowledgments

We wish to thank Michael Kunst for providing early access to the MERFISH data and expert insight and feedback on the data analysis. We also wish to thank Rohan Gala and Jonathan Ting for their comments on the initial draft of this work.

## Author contribution matrix

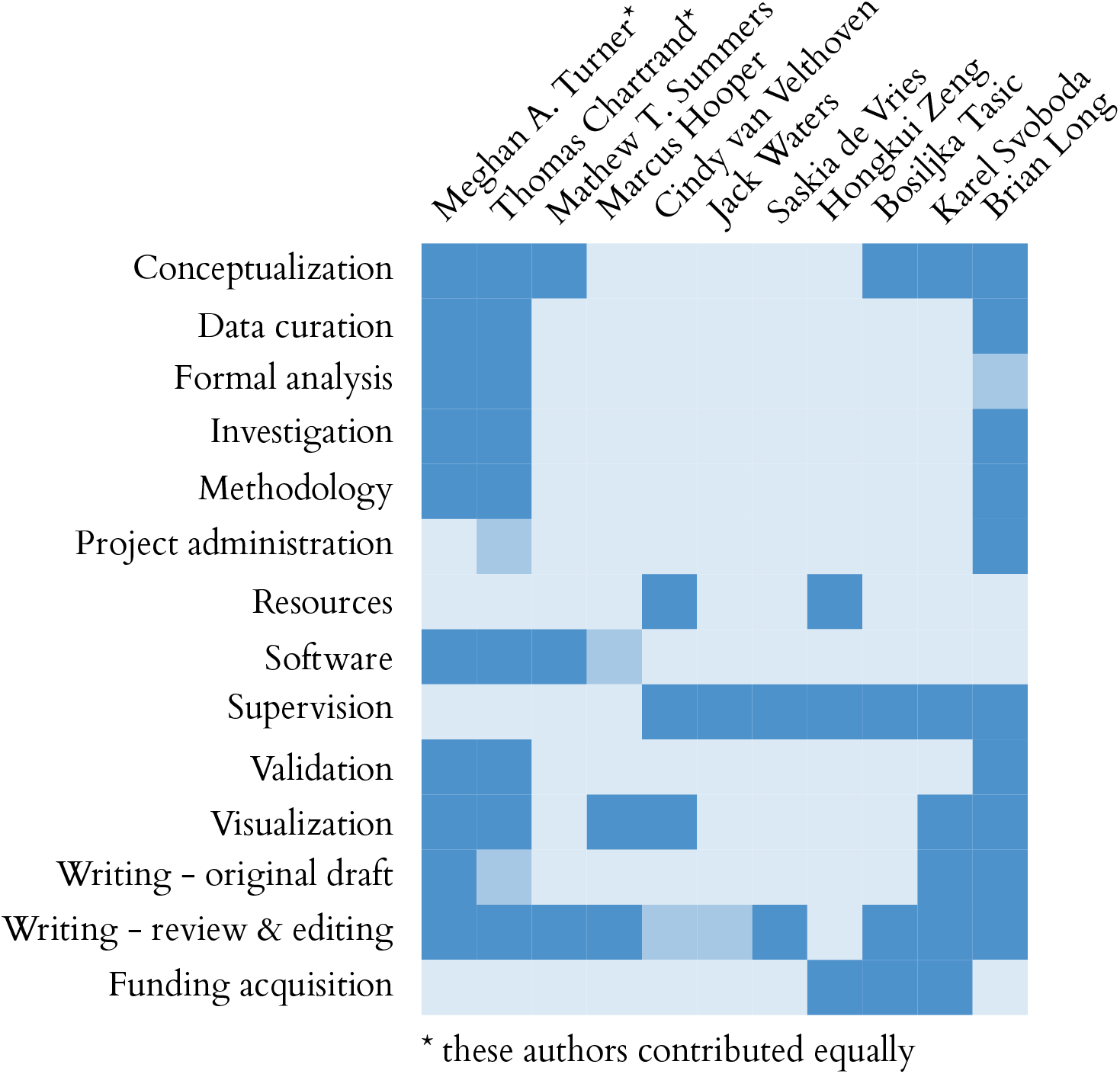

