## Supplemental Tables and Figures for "Exploring the correspondence between gene expression and thalamic nuclei using the THALMANAC resource"

**Table S1: Taxonomy of neuronal transcriptomic cell types *(t-types)* in the thalamus.** Thalamus subset of the ABC Atlas hierarchical taxonomy that are included in the THALMANAC resource. Preview of the first five rows is included in this manuscript; full table can viewed at table1_thalamus_taxonomy_CCN20230722.html.

| | neurotransmitter | class | subclass | supertype | cluster | n_cells_in_cluster |
|---|---|---|---|---|---|---|
| 0 | GABA | 12 HY GABA | 093 RT-ZI Gnb3 Gaba | 0431 RT-ZI Gnb3 Gaba_1 | 1580 RT-ZI Gnb3 Gaba_1 | 71 |
| 1 | GABA | 12 HY GABA | 093 RT-ZI Gnb3 Gaba | 0431 RT-ZI Gnb3 Gaba_1 | 1581 RT-ZI Gnb3 Gaba_1 | 128 |
| 2 | GABA | 12 HY GABA | 093 RT-ZI Gnb3 Gaba | 0431 RT-ZI Gnb3 Gaba_1 | 1582 RT-ZI Gnb3 Gaba_1 | 179 |
| 3 | GABA | 12 HY GABA | 093 RT-ZI Gnb3 Gaba | 0431 RT-ZI Gnb3 Gaba_1 | 1583 RT-ZI Gnb3 Gaba_1 | 78 |
| 4 | GABA | 12 HY GABA | 093 RT-ZI Gnb3 Gaba | 0431 RT-ZI Gnb3 Gaba_1 | 1584 RT-ZI Gnb3 Gaba_1 | 47 |

**Table S2: Glossary of thalamic nuclei.** Glossary of anatomical structure names and abbreviations in the thalamus as defined by the 3D Allen Mouse Brain CCFv3 and as published as part of the Allen Brain Cell (ABC) Atlas (see their parcellation annotation terms page: https://alleninstitute.github.io/abc_atlas_access/_static/Allen-CCF-2020/substructure.html). All analyses were done on the structure level of the ontology; substructures only provided for completeness.

| Division | Structure Abbreviation | Substructure Abbreviation | Substructure Name |
|---|---|---|---|
| TH | AD* | AD | Anterodorsal nucleus |
| TH | AM* | AMd | Anteromedial nucleus, dorsal part |
| TH | AM* | AMv | Anteromedial nucleus, ventral part |
| TH | AV* | AV | Anteroventral nucleus of thalamus |
| TH | CL* | CL | Central lateral nucleus of the thalamus |
| TH | CM* | CM | Central medial nucleus of the thalamus |
| TH | Eth | Eth | Ethmoid nucleus of the thalamus |
| TH | IAD* | IAD | Interanterodorsal nucleus of the thalamus |
| TH | IAM | IAM | Interanteromedial nucleus of the thalamus |
| TH | IGL | IGL | Intergeniculate leaflet of the lateral geniculate complex |
| TH | IMD* | IMD | Intermediodorsal nucleus of the thalamus |
| TH | IntG | IntG | Intermediate geniculate nucleus |
| TH | LD* | LD | Lateral dorsal nucleus of thalamus |
| TH | LGd* | LGd-co | Dorsal part of the lateral geniculate complex, core |
| TH | LGd* | LGd-ip | Dorsal part of the lateral geniculate complex, ipsilateral zone |
| TH | LGd* | LGd-sh | Dorsal part of the lateral geniculate complex, shell |
| TH | LGv | LGv | Ventral part of the lateral geniculate complex |
| TH | LH* | LH | Lateral habenula |
| TH | LP* | LP | Lateral posterior nucleus of the thalamus |
| TH | MD* | MD | Mediodorsal nucleus of thalamus |
| TH | MG | MGd | Medial geniculate complex, dorsal part |
| TH | MG | MGm | Medial geniculate complex, medial part |
| TH | MG | MGv | Medial geniculate complex, ventral part |
| TH | MH* | MH | Medial habenula |
| TH | PCN* | PCN | Paracentral nucleus |
| TH | PF* | PF | Parafascicular nucleus |
| TH | PIL | PIL | Posterior intralaminar thalamic nucleus |
| TH | PO* | PO | Posterior complex of the thalamus |
| TH | POL | POL | Posterior limiting nucleus of the thalamus |
| TH | PoT | PoT | Posterior triangular thalamic nucleus |
| TH | PP | PP | Peripeduncular nucleus |
| TH | PR | PR | Perireunensis nucleus |
| TH | PT | PT | Parataenial nucleus |
| TH | PVT* | PVT | Paraventricular nucleus of the thalamus |
| TH | RE* | RE | Nucleus of reuniens |
| TH | RH | RH | Rhomboid nucleus |
| TH | RT* | RT | Reticular nucleus of the thalamus |
| TH | SGN | SGN | Suprageniculate nucleus |
| TH | SMT | SMT | Submedial nucleus of the thalamus |
| TH | SPA* | SPA | Subparafascicular area |
| TH | SPFm | SPFm | Subparafascicular nucleus, magnocellular part |
| TH | SPFp | SPFp | Subparafascicular nucleus, parvicellular part |
| TH | SubG | SubG | Subgeniculate nucleus |
| TH | TH-unassigned | TH-unassigned | Thalamus, unassigned |
| TH | VAL* | VAL | Ventral anterior-lateral complex of the thalamus |
| TH | VM* | VM | Ventral medial nucleus of the thalamus |
| TH | VPL* | VPL | Ventral posterolateral nucleus of the thalamus |
| TH | VPLpc | VPLpc | Ventral posterolateral nucleus of the thalamus, parvicellular part |
| TH | VPM* | VPM | Ventral posteromedial nucleus of the thalamus |
| TH | VPMpc* | VPMpc | Ventral posteromedial nucleus of the thalamus, parvicellular part |
| TH | Xi | Xi | Xiphoid thalamic nucleus |
| HY | ZI | ZI-unassigned | Zona incerta, unassigned |

* Indicates this thalamic nucleus is highlighted in main figures.

**Table S3: Supplemental Data: Thalamic nuclei-to-cluster annotated matches.**
This supplemental data table can be accessed at: https://github.com/AllenNeuralDynamics/abc-merfish-analysis/blob/main/src/abc_merfish_analysis/resources/annotations_c2n_combined.csv

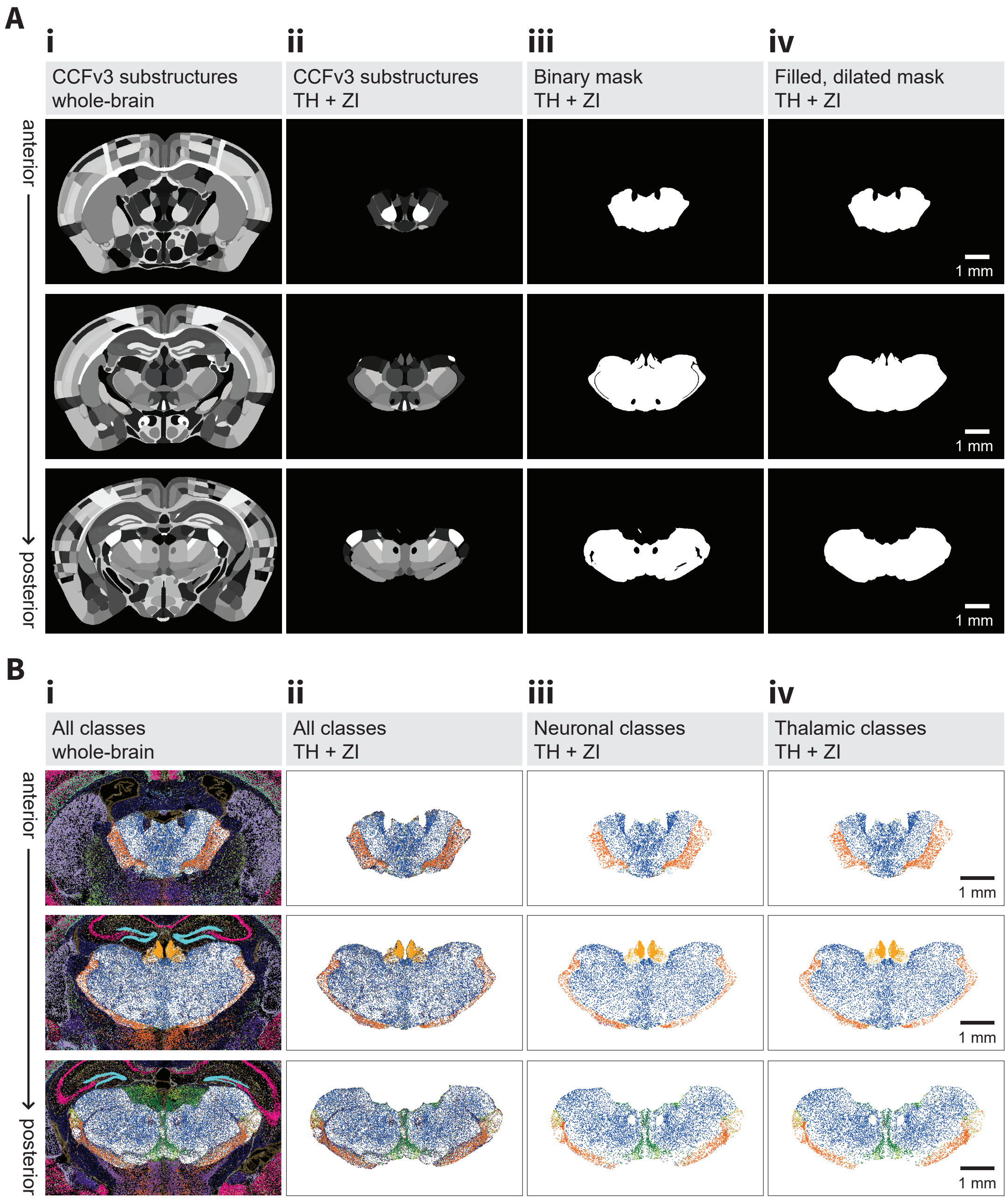


**Figure S1: Selection of the thalamic cells from the ABC Atlas mouse whole-brain MERFISH dataset.**
**(A)** Illustration of subsetting the ABC Atlas whole-brain dataset to a thalamic dataset based on spatial CCFv3 parcellation labels, illustrated in three example sections, from anterior (top row) to posterior (bottom row). Steps are performed in order from (i) to (iv):
(i) CCFv3 parcellation substructure labels, colored in greyscale, for the whole brain.
(ii) CCFv3 parcellation substructure labels for those substructures belonging to the thalamus (TH) and zona incerta (ZI).
(iii) Binary mask of the TH+ZI substructures from (ii).
(iv) Binary mask after inclusion of the interior white matter tracts (which appear as black holes in (iii)) and dilation of the outer boundary by 20um.
*Caption continues on next page.*

**Figure S1: Selection of the thalamic cells from the ABC Atlas mouse whole-brain MERFISH dataset.**
*Caption continued from previous page.*
**(B)** Workflow for subsetting the ABC Atlas to a thalamus-specific dataset based on scRNA-seq transcriptomic type class labels, illustrated in the same three example sections from (A). Steps are performed after those in (A), in order from (i) to (iv):
(i) All cells in the ABC Atlas whole brain dataset, colored by their class and overlaid on the filled, dilated TH+ZI binary mask from (A).
(ii) All cells that fall within the filled, dilated TH+ZI binary mask from (A).
(iii) All neuronal cells in the TH+ZI after filtering out cells belonging to the four non-neuronal classes (30 Astro-Epen, 31 OPC-Oligo, 33 Vascular, 34 Immune).
(iv) The final subset of TH+ZI cells used in this work, after filtering out of cells not belonging to the following neuronal classes: (12 HY GABA, 17 MH-LH Glut, 18 TH Glut, 19 MB Glut, and 20 MB GABA).

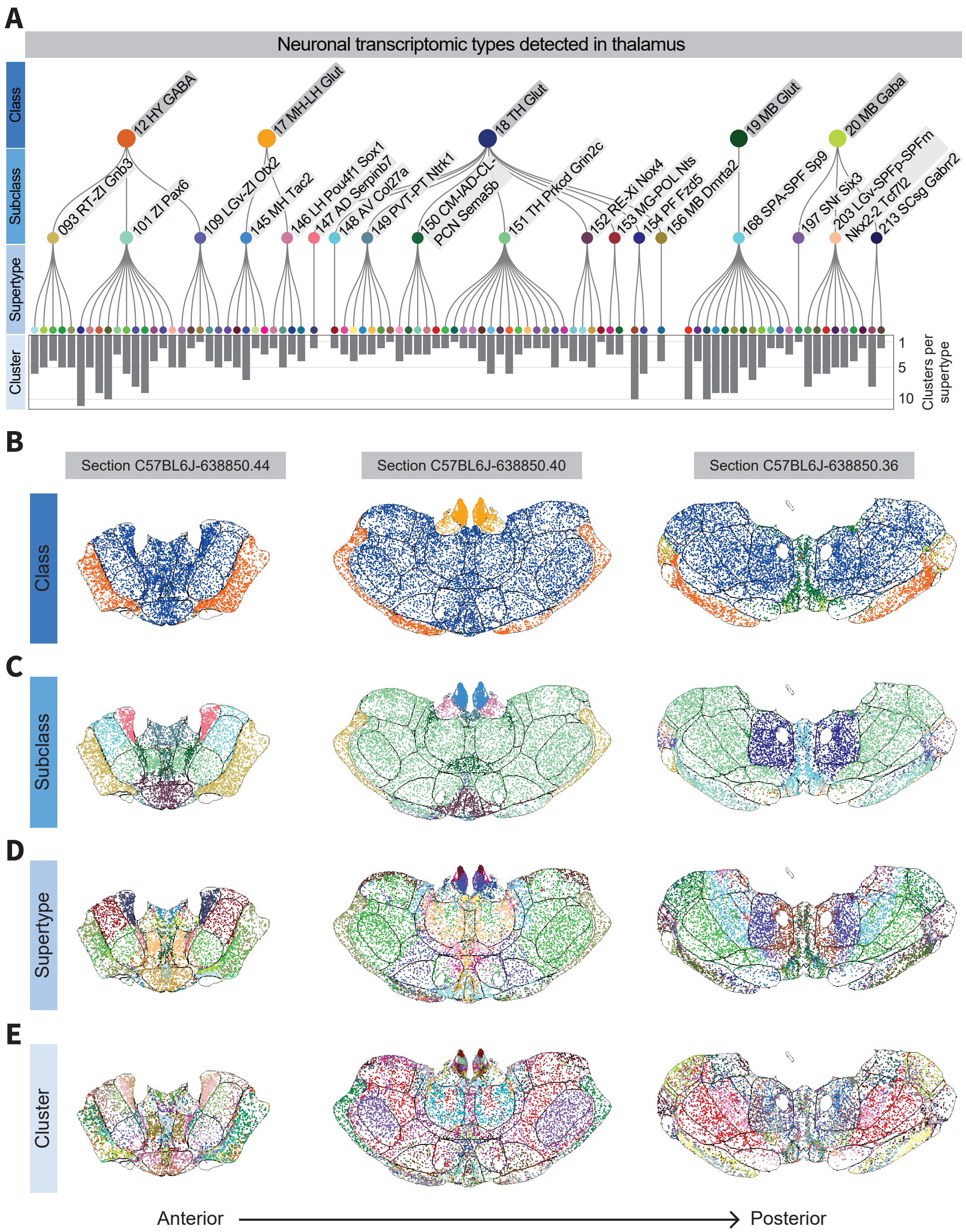


**Figure S2: Hierarchy of the transcriptomic taxonomy in the thalamus.**
**(A)** Dendrogram illustrating the hierarchical taxonomy organization of the neuronal transcriptomic types detected in the thalamus. **(B-E)** Spatial location of MERFISH cells in the thalamus (sections C57BL6J-638850.44, C57BL6J-638850.40, C57BL6J-638850.36) in each taxonomy level, from most to least broad: **(B)** class, **(C)** subclass, **(D)** supertype, **(E)** cluster.

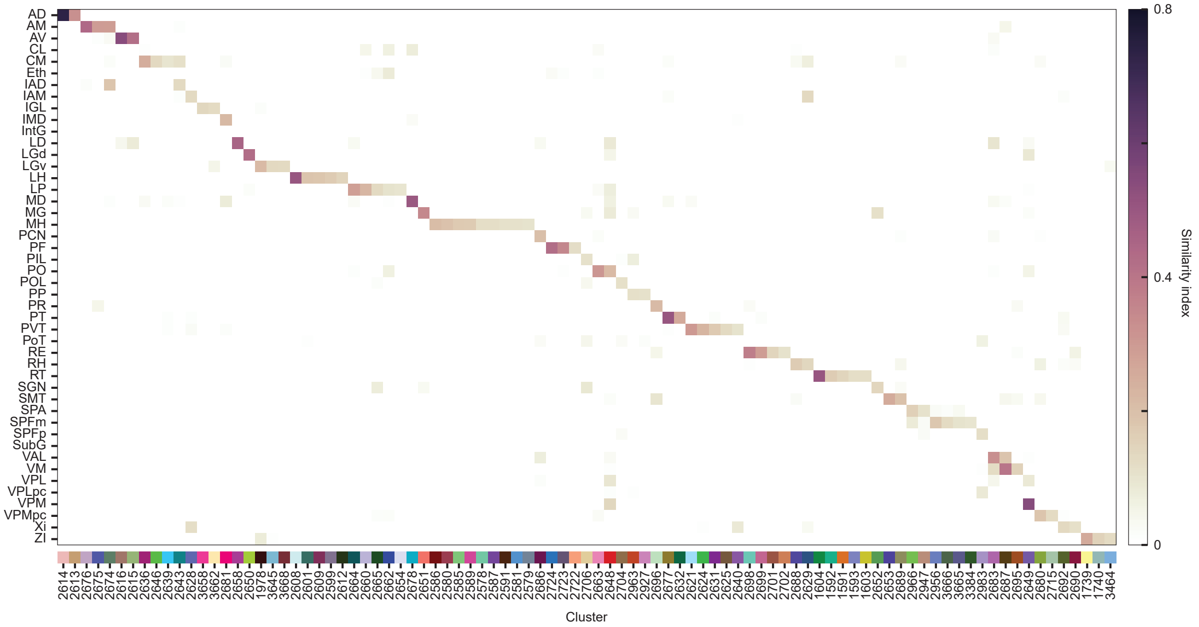


**Figure S3: Cell-wise similarity heatmaps for all thalamic nuclei.** Extended version of **Figure 2A** showing all thalamic nuclei on the x-axis. Heatmap quantifying the similarity of anatomically defined boundaries of thalamic nuclei to the spatial distribution of transcriptomic clusters from the scRNA-seq taxonomy. Thalamic nuclei names and abbreviations are listed in **Table S2**.

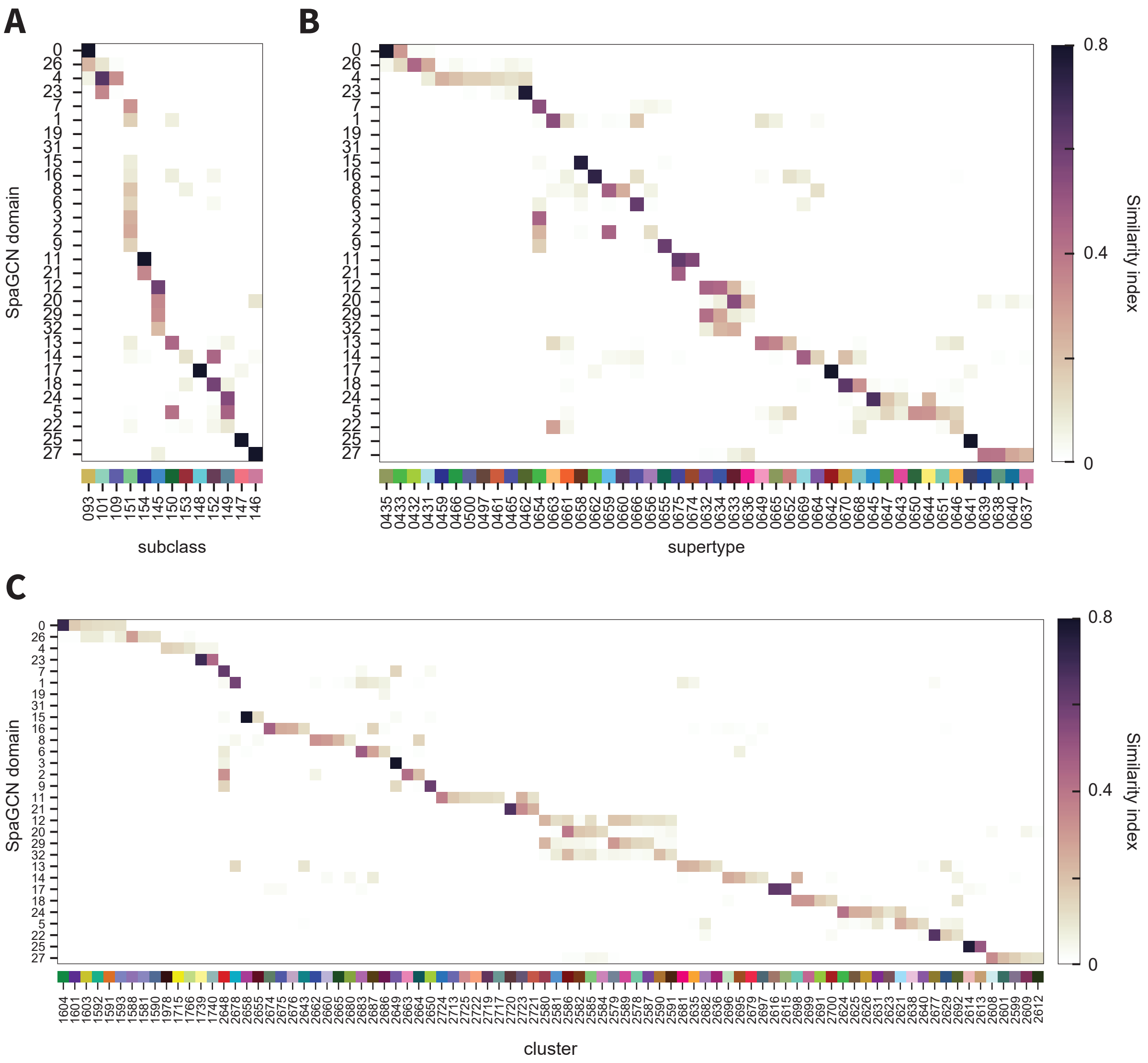


**Figure S4: Similarity of SpaGCN domains to transcriptomic cell types.**
Heatmaps quantifying the cell-wise similarity of SpaGCN domains to transcriptomic cell types, calculated with the Dice coefficient (see *Methods*), at three taxonomy levels: (A) subclasses, (B) supertypes, (C) clusters. SpaGCN domains are in the same order along the y-axis across all three panels.

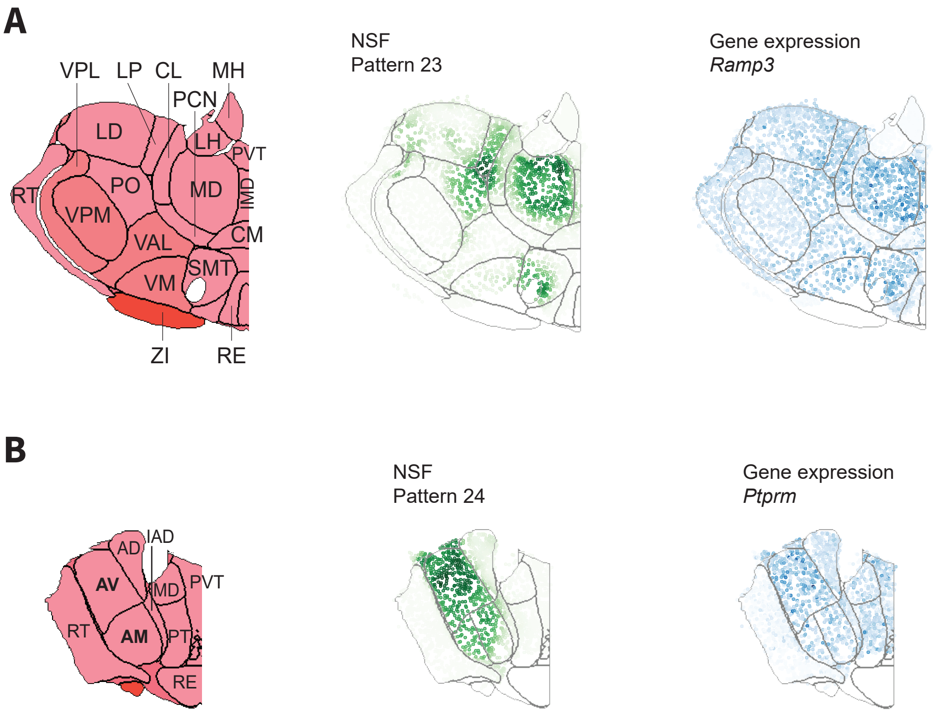


**Figure S5: Gene expression patterns that correspond to NSF patterns.**
The graded gene expressions extracted by NSF can be used to determine genes that show similar patterns: strong spatial structure, but in a graded manner that would be less significant when searching for differential expression between nuclei. **(A)** *Ramp3* shows the strongest differential contribution to NSF pattern 23, which is predominantly localized to MD, as well as other nuclei such as LP, PO, and SMT. **(B)** *Ptprm* shows the strongest differential contribution to NSF pattern 24, which is predominantly localized to AV and AM.
